# Differential causal involvement of human auditory and frontal cortices in vocal motor control

**DOI:** 10.1101/2020.06.08.139881

**Authors:** Araceli R. Cardenas, Roozbeh Behroozmand, Zsuzsanna Kocsis, Phillip E. Gander, Kirill V. Nourski, Christopher K. Kovach C, Kenji Ibayashi, Marco Pipoly, Hiroto Kawasaki, Matthew A. Howard, Jeremy D.W. Greenlee

## Abstract

Speech motor control requires integration of sensory and motor information. Bidirectional communication between frontal and auditory cortices is crucial for speech production, self-monitoring and motor control. We used cortical direct electrical stimulation (DES) to functionally dissect audio-motor interactions underlying speech production and motor control. Eleven neurosurgical patients performed a visually cued vocal task in which a short auditory feedback perturbation was introduced during vocalization. We evaluated the effect of DES on vocal initiation, voice fundamental frequency (F0) and feedback-dependent motor control. DES of frontal sites modulated vocal onset latencies. Stimulation of different inferior frontal gyrus sites elicited either shortening or prolongation of vocal latencies. DES distinctly modulated voice F0 at different vocalization stages. Frontal and temporal areas played an important role in setting voice F0 in the first 250 ms of an utterance, while Heschl’s gyrus was involved later when auditory input is available for self-monitoring. Vocal responses to pitch-shifted auditory feedback were mostly reduced by DES of non-core auditory cortices. Overall, we demonstrate that vocal planning and initiation are driven by frontal cortices, while feedback-dependent control relies predominantly on non-core auditory cortices. Our findings represent direct evidence of the role played by different auditory and frontal regions in vocal motor control.

## Introduction

Auditory feedback is necessary for speech motor control. Humans and non-human primates can rapidly adjust their voice acoustics in response to perceived changes in auditory input during vocalization (Eliades and Tsunada, 2018; Pomberger *et al.*, 2018). For example, the introduction of a pitch-shift in auditory feedback elicits “compensatory” vocal responses in humans (Behroozmand *et al.*, 2012) and other animals (Eliades and Tsunada, 2018).

Extant theories posit that a motor-to-sensory corollary signal represents the auditory feedback predicted according to the intended speech motor plan (Hickok *et al.*, 2011; Tian and Poeppel, 2014; Guenther, 2016). Auditory cortex suppression observed during vocalization is considered an indication of this process (Eliades and Wang, 2008; Flinker *et al.*, 2010; Greenlee *et al.*, 2011). During speech production the actual sensory input is compared to the expected one, as represented in this motor “efferent-copy”. Error detection during vocal self-monitoring triggers “compensatory” vocal responses. In tasks involving feedback-dependent speech motor control, activation of STG has been correlated with vocal performance (Chang *et al.*, 2013; Greenlee *et al.*, 2013; Behroozmand *et al.*, 2016). In consequence, speech self-monitoring and error detection are thought to rely on non-primary auditory cortices.

Accumulating evidence points to a frontal origin for the speech corollary signal (Schneider *et al.*, 2018; Eliades and Wang, 2019). Alexander *et al.* (1976) demonstrated in squirrel monkeys that electrical microstimulation of prefrontal areas was followed by suppression of single neuron activity in STG. Neurons in the primate frontal and premotor cortex not only exhibit pre-vocal activity (Hage and Nieder, 2013), but also respond to auditory stimuli (Hage and Nieder, 2015; Hage, 2018). These findings suggest that frontal areas can send input signals to auditory cortex and also receive outputs from it (Eliades and Wang, 2019). Along these lines, the consensus is that speech motor control relies on reciprocal interactions between auditory cortex in the temporal lobe and motor cortex in the frontal lobe. In humans, the arcuate and uncinate fasciculi likely enable bidirectional communication between these regions (Balezeau *et al.*, 2020). Still, the directionality of these interactions and their specific functional role in speech production are largely unknown.

In the last ten years, an increasing number of human studies reported the activation of speech motor cortices during purely listening conditions (Wilson *et al.*, 2004; Cheung *et al.*, 2016). Frontal motor areas not only encode auditory stimuli but also make use of auditory representations (Cogan *et al.*, 2014; Müsch *et al.*, 2020). On the one hand, these results support “mirror” models of speech, posing a common set of neural resources used in both perception and production. On the other hand, this “sensory” activation of motor circuitry challenges the functional relevance of audio-motor coactivations in speech generation.

In this context, causal approaches are necessary to dissect the contributions of auditory and motor areas to vocal production. Recently, Eliades *et al*. (2018) demonstrated that microstimulation of marmoset auditory cortex during vocalization evokes vocal responses similar to those elicited by pitch-shifting auditory feedback. Their results show that primate auditory cortex is causally involved in vocal self-monitoring. However, the correspondence between human and non-human primate auditory cortices is far from settled. In humans, electrical stimulation of HG is rarely possible and its effects on vocalization have been never assessed. Furthermore, there are not yet reports on the effect of motor areas stimulation on feedback-dependent vocal control, either in human or non-human primates. To disentangle the role of temporal and frontal cortices in human speech motor control, we applied cortical direct electrical stimulation (DES) at different hubs of the human auditory-motor network during vocalization. Particularly, we evaluated the effect of DES on vocal planning, initiation and sensorimotor control. We hypothesized that stimulation at higher levels of the audio-motor network (frontal areas and STG) could impact a putative corollary signal, required for vocal self-monitoring and error detection. In contrast, HG stimulation could affect vocal control by perturbing auditory feedback.

## Materials and Methods

Eleven neurosurgical patients were included in the study (**Supplementary Table 1**). Subjects were asked to produce and maintain a steady vocalization of the vowel /a/ as long as a visual cue (“GO”) was displayed. Their voice was captured with a microphone and fed back to the subject through insert earphones (**Fig. 1A**; ***See Supplementary methods***). In pitch-shift trials, phonation onset triggered the introduction of a short pitch-shift (+/−100 cents, 300ms duration) in auditory feedback after a jittered period of 1 ± 0.1s following voice onset. In a pseudo-randomized fashion bipolar DES was delivered to a selected pair of implanted ECoG contacts in half of the trials. DES was locked to the visual cue (3s duration; **Fig. (Boersma and Weenik, n.d.)1A**) and therefore started before vocalization. Vocal behavior was analyzed post-hoc (***See Supplementary methods***).

**FIGURE1.**
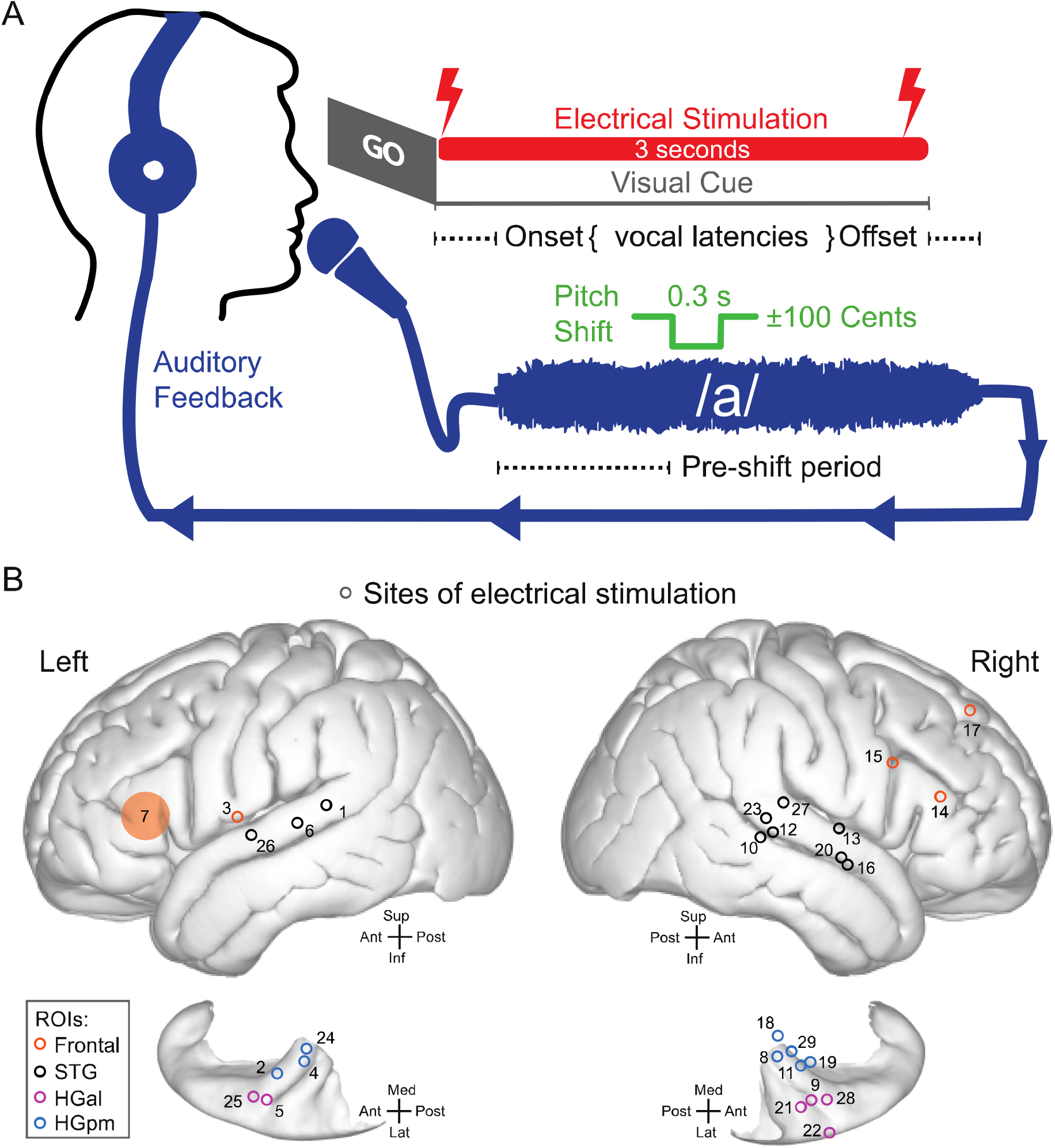
Task & DES contact locations. **Fig1A.** *Experimental Paradigm*. Subjects were asked to produce and maintain a steady vocalization of the vowel sound /a/ while a visual cue (“GO”) was displayed on a screen. Direct electrical stimulation (DES) was locked to the visual cue and persisted for 3 secs. In pitch-shift trials, the phonation onset triggered a brief perturbation (0.3 s) in the pitch of the auditory feedback (+/−100 Cents) after a jittered latency of 1±0.1 seconds. **Fig1B.** *Location of stimulation sites*. In pseudo-randomized trials, bipolar cortical electrical stimulation was delivered to a selected site pair of contacts in a block of approximately 40 trials. One pair of contacts was stimulated per block. Stimulation sites are depicted as empty circles midway between the locations of two contacts selected for bipolar stimulation. Circle colors denote ROI labelling for each DES site. Localization precision of site 7 is less than other sites, therefore, a larger shaded area is shown to account for this error.

We applied DES at 29 sites total, utilizing one pair of contacts per block and including both left and right hemispheres (**Fig. 1B, Supplementary Table 2**; sites indicated as the midpoint in MNI space between the utilized pair of contacts). Sites were grouped into 4 regions of interest (ROIs): posteromedial Heschl’s gyrus (HGpm, n=8), anterolateral Heschl’s gyrus (HGal, n=6), temporal (n=10) and frontal sites (n=5). Distinction of HGpm versus HGal was made based on responses to click train stimulation (Brugge *et al.*, 2009; Nourski *et al.*, 2016). Temporal sites included 9 STG sites and 1 site over superior temporal sulcus. In the frontal lobe, we stimulated 2 IFG sites, 1 IFS, 1 subcentral and 1 SFG site.

We investigated the effects of cortical stimulation on three aspects of vocal behavior. First, we used vocal response times to evaluate the impact of DES on vocal planning and initiation. Second, we quantified DES effect on voice fundamental frequency (F0) at the start of vocalization and during sustained phonation. Finally, to assess DES effect on feedback-dependent vocal control, we evaluated F0 responses to a brief pitch-shift in auditory feedback.

## Results

### Vocal initiation and termination are modulated by frontal DES

Vocal onset latencies were calculated as the time between the display of the visual go cue and vocalization onsets. DES at four of five frontal sites significantly affected vocal onset latencies [**Fig. 2**, Mann-Whitney U-test, p≤0.05]. These four sites included left IFG (L311, s7), right IFG (R320, s14), right SFG (R329, s17), and left subcentral gyrus (L275, s3; see also **Fig. 1B**). Interestingly, frontal DES resulted in either prolongation or shortening of vocal onset latencies. For example, stimulation of left IFG slowed vocal responses (L311, s7), while stimulation of right IFG resulted in significantly faster vocal responses (R320, s14). The only frontal site where DES did not affect vocal onset times (R322, s15) was located on the inferior frontal sulcus (**Fig. 1B**).

**FIGURE2.**
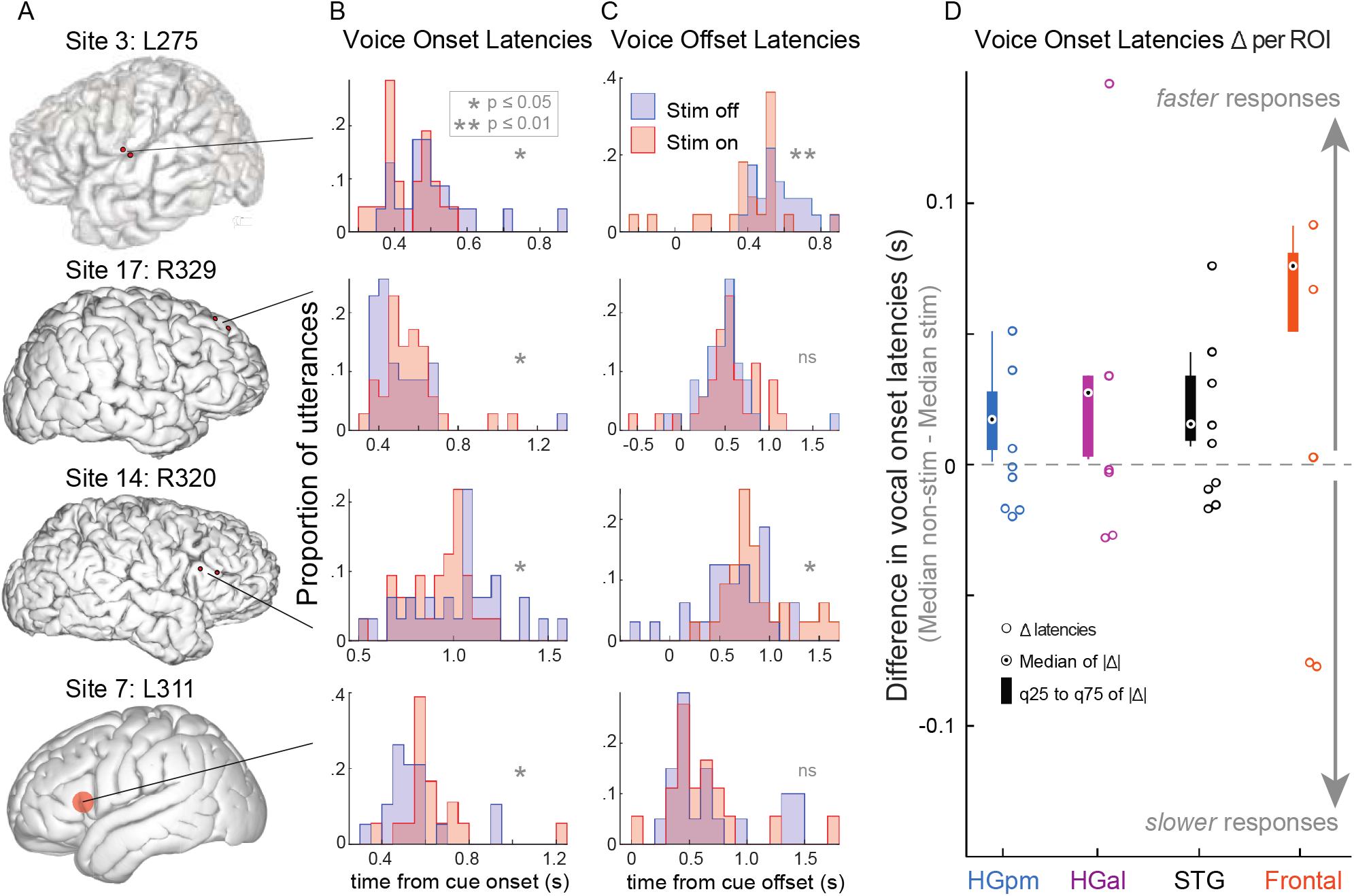
Effect of DES on vocal onset and offset times. **Fig2A-C**. ***DES on individual prefrontal sites affecting vocal response times.*** **Fig2A**. Location of each of stimulated site. **Fig2B**. Distributions of vocal onset latencies in stimulation (red) and non-stimulation (blue) trials. **Fig2C** Distributions for vocal offset latencies. Vocal offset latencies are negative when a subject stopped vocalizing before cue disappearance. Differences between distributions were tested with a Mann-Whitney U-test; the p level in each block is represented according to standard nomenclature (*p≤0.05, **p≤0.001). **Fig2D**. ***Change in vocal onset latencies elicited by DES on different cortical areas.*** The difference between median vocal onset times in non-stimulation and stimulation trials is marked with an open circle for each stimulation site and color coded for the different cortical ROIs. Values larger than zero indicate that DES shortened vocal onset times and values smaller than zero correspond to a prolongation in vocal latencies by DES. The absolute difference in response times, per contact, was used to evaluate DES effect regardless its direction (+/−). Box plots per area depict the median of these absolute values (pointed circles) from percentile 25 to 75.

We quantified the change in vocal response times evoked by DES by subtracting the median response times in *stimulation* trials from those in *non-stimulation* trials and grouped according to stimulated ROI (**Fig. 2D).** While stimulation of frontal sites (4/5) elicited significant effects, neither STG (0/10) nor HG (0/14) DES altered vocal latencies. To compare ROIs, we estimated the effect of DES on vocal latencies irrespective of its direction (i.e. prolongation vs. shortening) using the absolute value of DES-induced latency differences. Box plots in **Fig2D** represent the absolute change in vocal onset latencies per region. In comparison to other areas, frontal DES evoked larger magnitude latency effects than other ROIs (frontal 0.076s [0.051-0.081] vs. other ROIs 0.017s [0.007-0.033], t-test p≤0.05). These results offer evidence that frontal activity is crucial in vocal planning and initiation.

We also assessed the effect of DES on vocal *offset* latencies, defined as the time from cue offset to the end of vocalization. Vocal offset latencies were negative when subjects stopped vocalizing before cue disappearance. DES on two frontal sites (s3 [left subcentral] and s14 [right IFG], **Fig. 1B**) significantly modulated vocalization offsets (**Fig. 2C**, Mann-Whitney U-test p≤0.05). We did not find evidence that stimulation of other cortical sites influenced offset latencies.

### The setting of voice F0 at the start of an utterance is predominantly driven by frontal and STG cortices

We investigated the role of frontal and auditory areas in the setting and maintenance of voice F0. We analyzed F0 before the introduction of any perturbation in the auditory feedback. This pre-pitch-shift period (**Fig. 1A**) comprised 800 ms after voice onset. **Fig. 3A-D** depicts the effect of an exemplar site DES (s3) on voice F0 (see also **Supplementary Fig1)**. **Figure 3** shows individual trial voice F0 (Hz) traces for non-stimulation (**Fig. 3A**) and stimulation (**Fig3B**) trials. DES modulated both F0 level (**Fig. 3C**) and its trial-by-trial variability over the experiment (**Fig3D**). For each site, the change in F0 elicited by DES was calculated per time point (t) using the formula

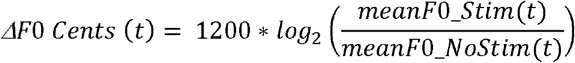

where the sign of the result indicates the direction of the change (**Fig. 3E**).

**FIGURE3.**
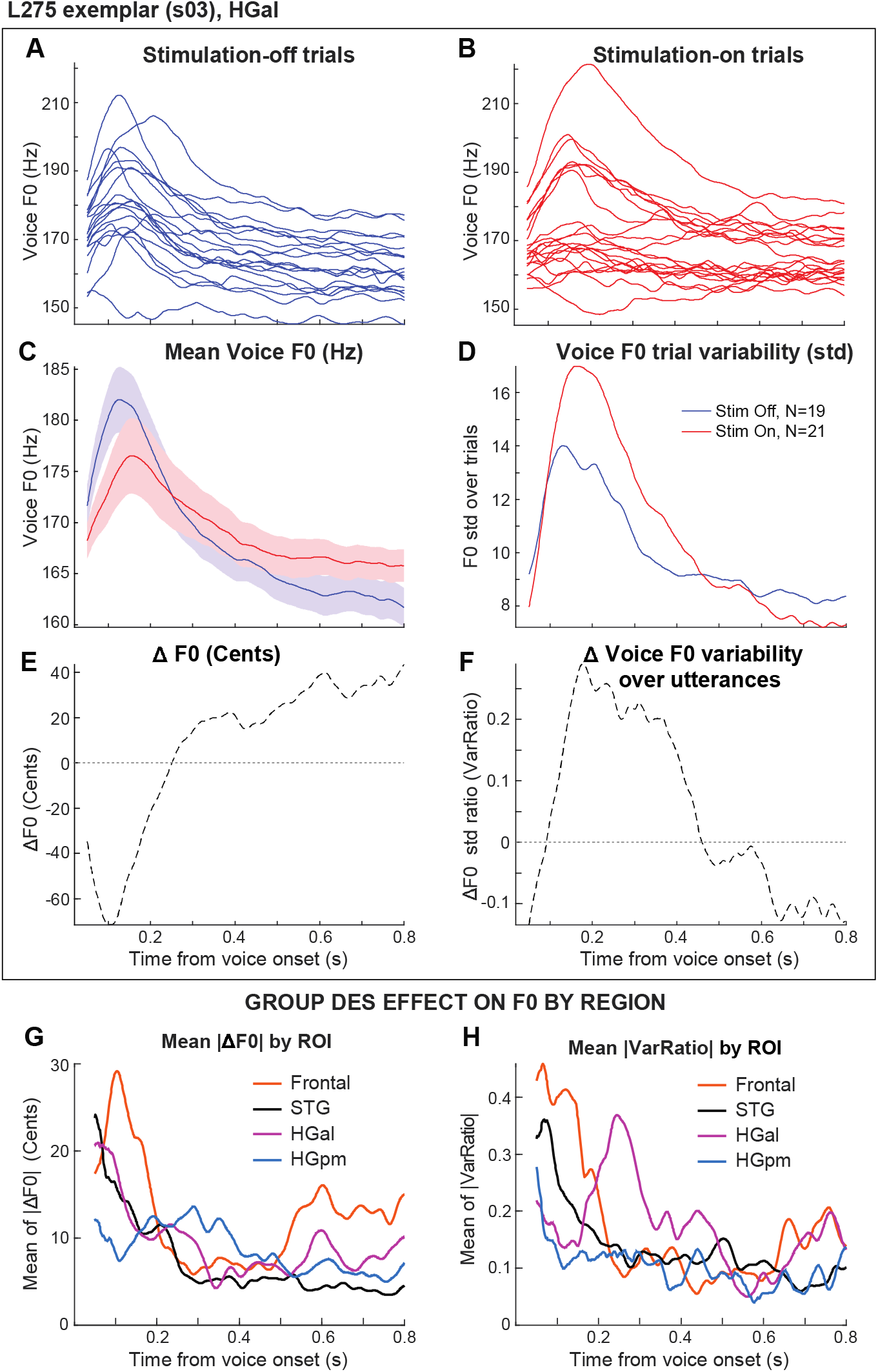
Effect of DES on voice F0 level and variability over utterances. **Fig3A-F**. *Effect of DES on voice F0 in an exemplary block*. **Fig3A-B** show single-utterance voice F0 traces in non-stimulation (blue) and stimulation trials (red) during the block in which frontal s03 was stimulated. F0 traces during the pre-shift period are aligned to voice onset. **Fig3C** shows mean F0 traces in non-stimulation and stimulation trials; shaded areas depict standard error over trials. **Fig3E** The change in F0 (ΔF0) elicited by DES was quantified in Cents using as reference the mean F0 in non-stimulation trials (dotted line, *See Methods*). Values larger than zero indicate that DES increased F0 and values smaller than zero correspond to a decrease in F0 evoked by DES. **Fig3D** depicts the standard deviation across non-stimulation (blue) and stimulation trials (red) in the pre-shift period. **Fig3F** DES-induced change in F0 variability over trials/utterances was quantified as a ratio (*VarRatio*) with respect to F0 standard deviation in non-stimulation trials (*See Methods*). Dotted line depicts the value of *VarRatio* during the pre-shift vocalization period in this block. Values larger than zero indicate that DES increased F0 variability and values smaller than zero correspond to a decrease in F0 variability evoked by DES. **Fig3G-H**. *Mean effec of DES on F0 level and inter-utterance variability per ROI*. To quantify the effect of DES per ROI independently of its direction, we averaged the absolute value of DES-induced changes (|ΔF0)| or |VarRatio|) over the sites of a ROI. **Fig3G** displays the average (absolute) effect of stimulation on F0 level (|ΔF0|) per ROI. **Fig3H** depicts the mean effect of stimulation on F0 variability (quantified as |VarRatio|) over trials per ROI.

We averaged DES-induced ΔF0 over sites to obtain estimates of mean change in F0 evoked by each ROI. Overall, stimulation of frontal sites and HGal lowered F0 early in the utterance (first 200 ms), while STG stimulation increased F0 in that period (**Supplementary Fig. 2A**). Stimulation of primary auditory cortex (HGpm) affected voice F0 only after this initial period (i.e. 200–500 ms after voice onset; HGpm 5.99 ± 3.61 vs. other ROIs −0.449 ± 1.42, t-test p≤0.05).

To obtain estimates of DES F0 effect regardless of effect *direction*, we also averaged the absolute change functions per ROI (**Fig. 3G**). This analysis showed that in the first 200 ms of an utterance, F0 was most modulated by frontal stimulation (frontal 21.87±7.78 vs. other ROIs 13.07±1.28 cents, t-test p=0.055), followed by STG and HGal.

As shown in **Fig. 3D** DES not only modulated voice F0, but also F0 variability (standard deviation) over utterances. DES alterations in F0 variability per time point (t) were quantified using a ratio index (VarRatio):

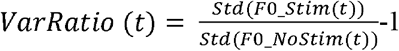

where the sign indicates the direction of the change evoked by DES (**Fig. 3F**). We averaged VarRatio values across sites in the same ROI. We found frontal DES increased F0 variability across utterances early in the vocalization (first 200 ms; 0.37±0.14). In contrast, HGal DES increased F0 variability later in an utterance (i.e. 200–500 ms after voice onset; **Fig. 3H**, **see also** **Supplementary Fig2. B**) (HGal 0.22±0.1 vs. other ROIs 0.11±0.01, t-test p≤0.05).

Taken together, these findings suggest that higher-order cortices (i.e. frontal and STG) set and modulate voice F0 very early in an utterance. HG seems to contribute mostly later, when auditory feedback is available for vocal self-monitoring.

### Voice F0 feedback-dependent responses rely primarily on non-core auditory cortices

We utilized an auditory feedback perturbation paradigm (i.e. pitch-shifted feedback, +/−100 cents) to examine the impact of DES on feedback-dependent vocal motor control. Voice F0 responses in the 400 ms following pitch-shifts were compared with and without stimulation. **Fig4. A** shows mean voice F0 traces in non-stimulation and stimulation trials for four different DES sites. The peak of the mean F0 response to the shift was identified (black arrows, **Fig. 4A**; ***see Supplementary Methods***) and quantified for comparison in non-stimulation and stimulation trials (**Fig. 4B)**.

**FIGURE4.**
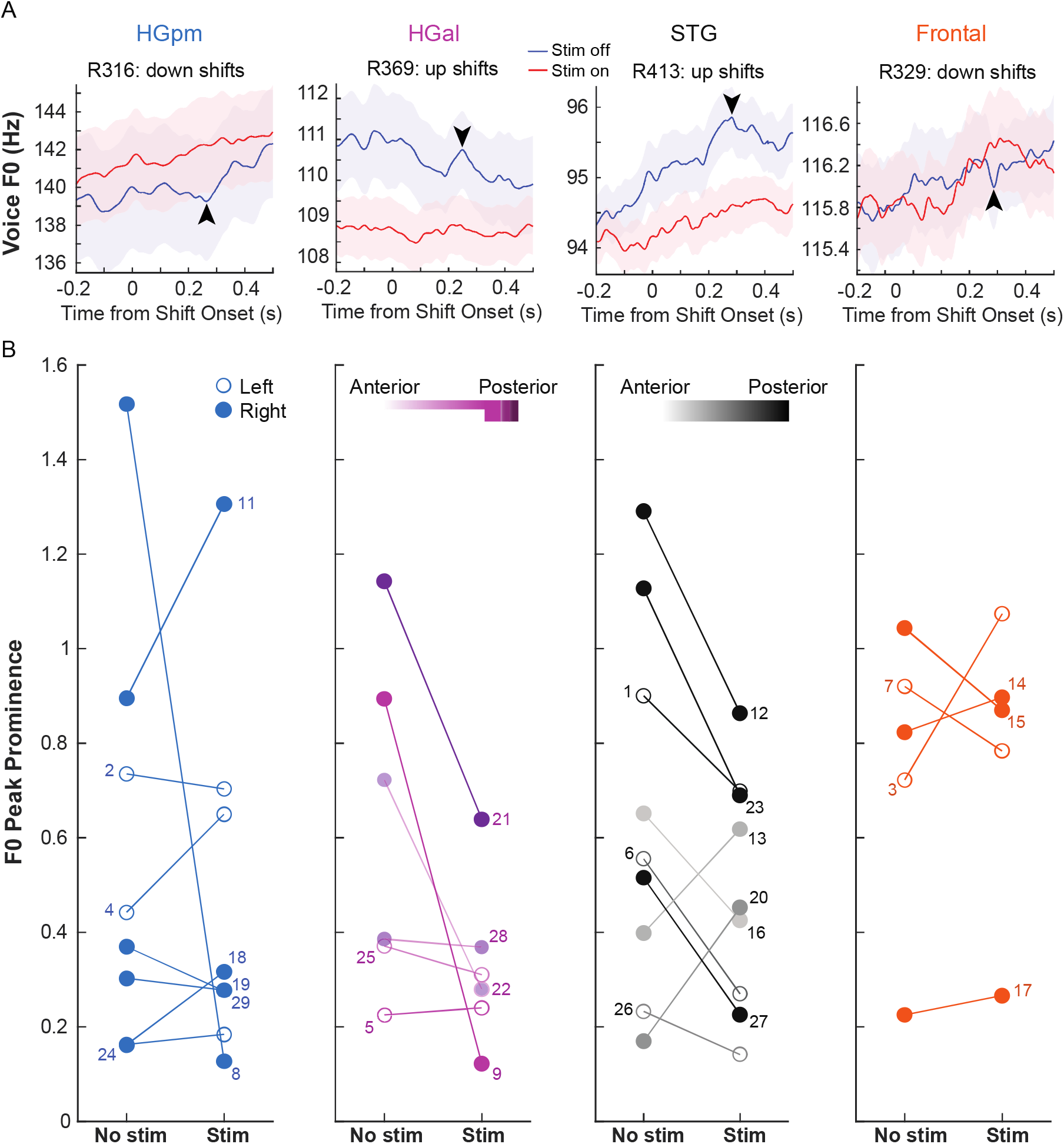
Effect of DES on feedback-dependent vocal control. **Fig4A**. ***Individual block examples per cortical area.*** The most prominent F0 peak in the response period was identified in detrended stimulation and non-stimulation average F0 traces. Notice that these traces are not normalized to any baseline period. Shade area represents standard error over utterances. Identified peaks are marked by an arrow. **Fig4B**. **Effects of DES on F0 peak responses per cortical area.** For each stimulated site, F0 response peak prominence in non-stimulation and stimulation trials is shown. For HGal and STG the antero-posterior location of sites is color coded.

The prominence of post-shift F0 peak responses decreased most commonly and noticeably during HGal stimulation followed by STG DES (**Fig. 4B**). Electrical stimulation of all sites in the posterior STG reduced the prominence of the peak F0 response to pitch-shifted auditory feedback (**Fig. 4B**, **see also** **Fig. 1B**). The only two temporal sites which did not follow this pattern were located more anteriorly in the middle STG (s13, s20 **Fig. 4B**, **see also** **Fig. 1B**). In the case of HGal, stimulation of more posteromedial sites evoked larger decreases in peak prominence. These results show that non-core auditory cortices play a crucial role in feedback-dependent vocal motor control. Interestingly, stimulation of *right* HGal (filled circles, **Fig. 4B**) and posterior STG appeared to reduce F0 peak responses to a greater degree than *left* side stimulation (open circles, **Fig. 4B**).

## Discussion

By reversibly perturbing cortical function through DES in the human brain we have shown that frontal and auditory cortices make differential contributions to vocal motor control. In agreement with the predominantly motor role of frontal areas in primate vocalization (Gavrilov *et al.*, 2017), we found that frontal DES affected vocal initiation as indexed by vocal onset latencies. Extant speech production models posit the involvement of supplementary motor area (SMA), pre-SMA, IFS, and PMC in speech planning (Hickok and Poeppel, 2007; Hickok *et al.*, 2011; Tourville and Guenther, 2011). Here we have presented supporting exemplars of DES on three of those areas.

Interestingly, stimulation of different IFG sites either prolonged or shortened vocal initiation in response to a visual cue. The bi-directionality of this effect is consistent with previous reports of IFG activity modulating speech timing. For example, applying direct cortical cooling to IFG during connected speech (Long *et al.*, 2016) evoked speech speeding or slowing once it had been initiated. In contrast to that study, our task did not involve articulatory motor activity, but the sustained phonation of a simple speech token in response to a visual cue. In principle, our results offer causal evidence of IFG role in vocal planning. They also suggest that the effects of IFG stimulation in speech timing might not be fully explained by its contributions to vocal articulation. The documented role of IFG in executive function could well account for the effects we observed. In this sense, our findings could support the view that Broca’s area is a functionally heterogeneous region, comprising a domain-general “multiple-demand network” (Fedorenko and Blank, 2020).

Most accounts of vocal production include some sort of “target” representation of the expected vocal output. However, produced vocalizations differ from that representation and from each other. The importance of “trial-by-trial” variation in vocal production is recognized in models of vocal control and learning (Garst-Orozco *et al.*, 2014; Parrell *et al.*, 2019). In songbirds, vocal variability has a central origin (Sober *et al.*, 2008). Here we offer a first account of the neural sources of F0 variability in primates. Particularly, we showed that different cortices modulate not only F0 level, but also its variability across vocalizations, and that this modulation is time-specific. For example, frontal DES changed F0 and increased F0 variability across trials in the first 250 ms of vocalization. These changes suggest a causal role for frontal cortex in setting or achieving an F0 target at the start of vocalization. In contrast to frontal and STG sites, HG sites seem to modulate F0 only later, when auditory feedback is likely utilized for self-monitoring.

Finally, we found that feedback-dependent vocal control was maximally disrupted by DES of posterior STG and HGal. Specifically, the peak F0 response to a pitch-shift in auditory feedback was more consistently reduced or abolished by stimulation of non-core auditory cortices. This observation causally demonstrates that sensorimotor vocal control in humans relies on HGal and STG. The dominant role of STG in feedback-dependent motor control contrasts with its limited contribution to maintaining F0 during sustained phonation. This observation suggests that STG may not modulate F0 *per se* but could be specifically involved in error detection.

In our study, DES of both right HGal and right posterior STG appear to have greater influence on vocal responses than the left-sided cohorts. Our observation is congruent with the report by Eliades et al. (2018) that in two marmosets right-sided electrical microstimulation of auditory cortex evoked stronger changes in vocal pitch (Eliades and Tsunada, 2018). Given that the right hemisphere is overrepresented in the present study, our results warrant additional research to examine hemispheric contributions.

Our findings highlight how direct causal approaches can be used to evaluate the comparative contributions of brain areas to a particular behavior. Notably, the number of trial repetitions per site and the duration of electrical stimulation in surgical patients are restricted by clinical and practical considerations. As a result of these constraints, our coverage of frontal regions was varied and sparse, limiting our insights into the role of specific frontal subregions in vocal motor control. Nevertheless, our findings represent the first effort to directly affect different hubs of the human audio-motor network during speech production and sensorimotor control.

## Abbreviations

DES: (cortical direct electrical stimulation)
ECoG: (electrocorticography)
F0: (voice fundamental frequency)
HGal: (anterolateral Heschl’s gyrus)
HGpm: (posteromedial Heschl’s gyrus)
IFG: (inferior frontal gyrus)
IFS: (inferior frontal sulcus)
ROI: (region of interest)
SFG: (superior frontal gyrus)
STG: (superior temporal gyrus)

## Acknowledgements

We primarily acknowledge the generosity of our patients. We also thank our colleagues in the Human Brain Research Laboratory for their feedback. We specially appreciate the assistance of Haiming Chen and Beau Snoad during the experiments.

## Funding Information

This work was supported by NIH grants DC015260 and DC004290.

## Competing Interests

The authors have no conflicts of interest to declare relevant to this work.

## SUPPLEMENTARY MATERIAL

**Supplementary Table 1.** Subject characteristics.

| DES side<sup>1/</sup><br>subject | Age | Sex<sup>2</sup> | Language<br>dominance<sup>3</sup> | Handedness | Number of DES sites |  |  |  |
| --- | --- | --- | --- | --- | --- | --- | --- | --- |
|  |  |  |  |  | HGpm | HGal | Temporal | Frontal |
| **L275** | 20 | M | L | +90 | 1 | 0 | 1 | 1 |
| **L307** | 20 | M | L | +100 | 1 | 1 | 1 | 0 |
| **L311** | 66 | F | L | R | 0 | 0 | 0 | 1 |
| **R316** | 32 | F | - | +100 | 1 | 1 | 1 | 0 |
| **R320** | 51 | F | - | +100 | 1 | 0 | 2 | 1 |
| **R322** | 28 | F | - | +100 | 0 | 0 | 1 | 1 |
| **R329** | 20 | M | - | +100 | 0 | 0 | 0 | 1 |
| **R335** | 33 | M | L | +75 | 2 | 0 | 1 | 0 |
| **R369** | 29 | M | L | +100 | 0 | 2 | 1 | 0 |
| **L403** | 56 | F | L | +100 | 1 | 1 | 1 | 0 |
| **R413** | 22 | M | R | -100 | 1 | 1 | 1 | 0 |
|  |  |  |  | **Total** | **8** | **6** | **10** | **5** |

<sup>1</sup>L = left; R = right.
<sup>2</sup>F = female, M = male

**Supplementary Table 2.**
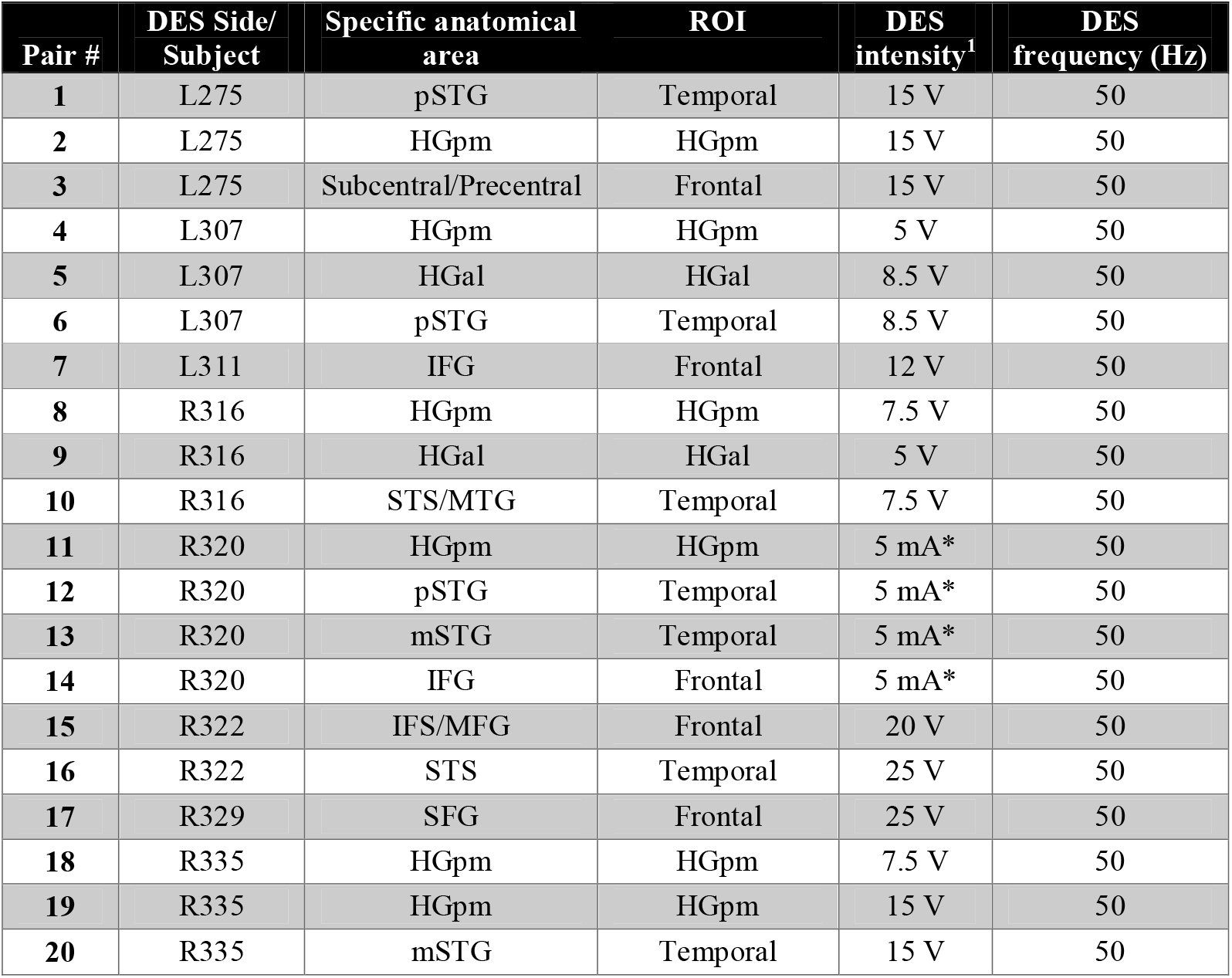
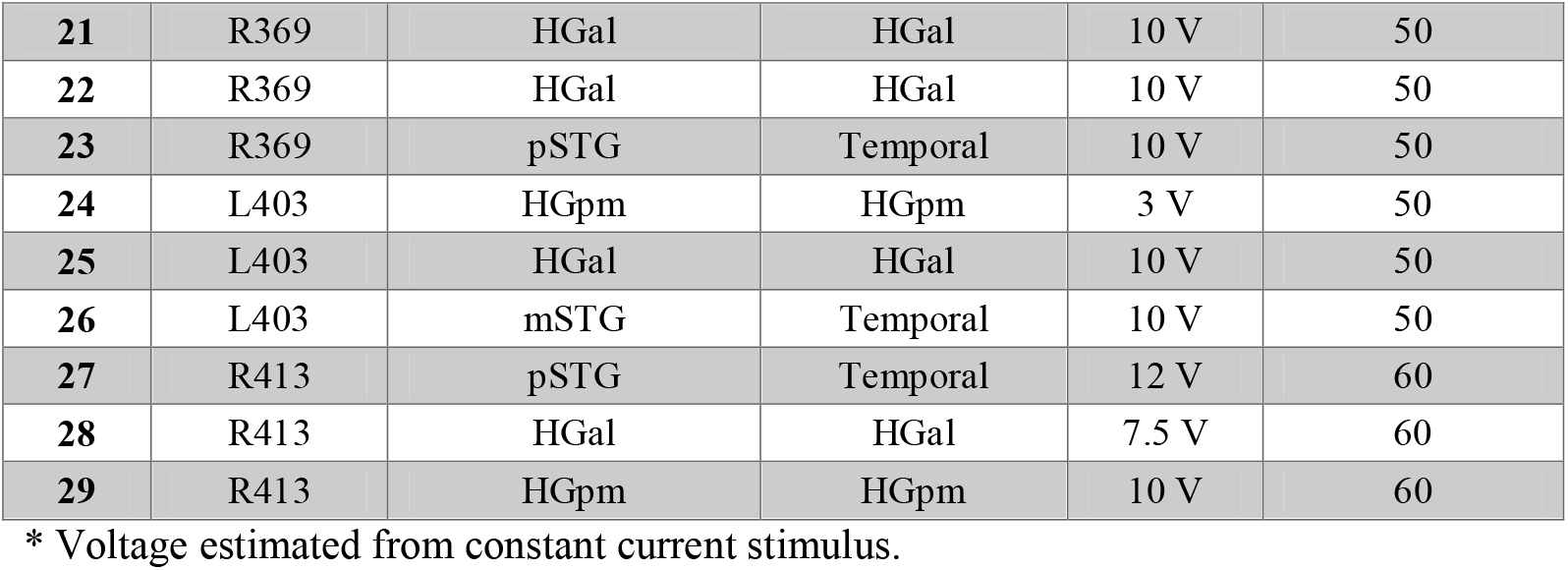
DES sites and stimulation parameters.

## SUPLEMENTARY METHODS

### Subjects

Data here presented was collected from 11 neurosurgical patients (5 females, 6 males, age 20-66 years old, median age 29 years old; See **Supplementary Table 1**). We included the data from blocks in which electrical stimulation at 50-60 Hz was applied to any region of interest (ROI) during the task. Ten subjects underwent surgical placement of subdural electrodes for seizure onset zone localization. These subjects completed one block of the task, without electrical stimulation, days prior to the DES session. One subject (L311) performed the task in the operating room during awake craniotomy for tumor resection. No hearing or language impairments were detected in any of the subjects. Research protocols were approved by the University of Iowa Institutional Review Board and the National Institutes of Health. Written informed consent was obtained from all subjects. Research participation did not interfere with acquisition of clinically required data, and subjects could rescind consent at any time without interrupting their clinical evaluation.

### Cortical direct electrical stimulation

**Supplementary Table 2** shows the electrical stimulation parameters used per site. Stimulation site selection was informed by the results a first run of the task without electrical stimulation. For each subject, we identified the pitch-shift direction (+/−100 cents) that elicited larger vocal responses. Neural data from that first session was analyzed and contacts with selective high gamma power responses were identified. ‘Selective’ sites were those showing differential responses in speaking versus listening (i.e. playback) trials, and in pitch-shifted versus control trials (Chang *et al.*, 2013; Greenlee *et al.*, 2013; Behroozmand *et al.*, 2016). Site selection was also constrained by clinical considerations including available coverage of HG, STG, and frontal cortices. Because L311 was tested in the operating room, a baseline non-stimulation pitch-shift block was not possible, and the site of DES was chosen based empirically on anatomic landmarks (i.e., IFG). Distinction of HGpm versus HGal sites was made based on responses to click train stimulation; particularly, we evaluated the presence of short-latency peaks (<20 ms) and frequency-following responses in the averaged auditory evoked potentials [see(Brugge *et al.*, 2009; Nourski *et al.*, 2016). HGpm was interpreted as putative core (primary and primary-like) auditory cortex while HGal was considered non-core auditory cortex.

Electrical stimulation was delivered using constant-voltage (n=10 subjects, Grass SD-9, Natus Neurology, Inc., Warwick, RI) and constant-current (n=1, Tucker-Davis Technologies) stimulators. We delivered trains (50Hz [n=10 subjects] or 60Hz [n=1 subject]) of 0.2ms duration biphasic, charge-balanced pulses of varying intensity (**Supplementary Table 2**). Stimulus intensity in all cases was chosen to be below after-discharge thresholds for each site as determined by continuous ECoG recordings over cortical areas adjacent to stimulation sites. The ECoG recordings were monitored by epilepsy monitoring unit staff for all ten subjects tested there. The recordings from subject L311 were monitored by an epilepsy neurologist in the operating room; none of these personnel were affiliated with this study.

Subjects were asked about any stimulus-induced percepts or self-reported or experimenter-observed speech production or perception difficulties, and if any were evident, stimulus intensity was lowered to avoid these. This testing was completed with test stimulation epochs at each site prior to beginning the DES experimental block.

Electrical stimuli were delivered to pairs of the clinical ECoG contacts (n=10 subjects, Ad-Tech, Inc, Oak Creek, WI). These included flat disc electrodes embedded in silastic grid arrays for frontal and STG sites, and penetrating depth electrode arrays of macrocontact ring contacts for HG sites. Stimulation for L311 was delivered using a retractor-fixed silver ball tip probe on the IFG surface in the operating room.

### Anatomical Reconstructions

A high-resolution T1-weighted structural MRI of the brain was acquired for each subject before and after electrode implantation. In three subjects group images were acquired from a 3T Siemens TIM Trio scanner with a 12-channel head coil (MPRAGE: 0.78 × 0.78 mm, slice thickness 1.0 mm, TR = 2.530 s, TE = 3.520 ms, average of two). In a second group of 7 subjects, images were acquired from a 3T GE Discovery MR750w scanner with a 32-channel head coil (IR-FSPGR: 1 × 1 mm, slice thickness 0.8 mm, TR = 8.504 ms, TE = 3.288 ms). To determine the location of lateral temporal surface recording electrode contacts on the preoperative structural MRI, these images were coregistered to post-implantation structural MRIs using a 3D linear registration algorithm (Functional MRI of the Brain Linear Image Registration Tool) and custom Matlab v.9.0 (MathWorks, Natick, MA) scripts using guidance from computed tomography (CT) scans (in-plane resolution 0.51 × 0.51 mm, slice thickness 1.0 mm). When possible, results were also compared with intraoperative photographs to ensure reconstruction accuracy. In the subject performing the task intraoperatively (S311), contact location was reconstructed from available image studies and experimental records. In this case, low quality contrasted pre-surgical images limited the detail of brain surface reconstruction. In consequence, we used a 3D MNI template to exemplify stimulation site location.

### Voice recordings and data analysis

Voice sound was captured by a microphone (Beta 87C, Shure) located near the subject’s mouth, amplified (10 dB gain; Ultralite MK3, MOTU), and passed through a harmonizer (Eclipse, Eventide), so a pitch shift of +100/−100 cents was introduced to the auditory feedback. The voice and feedback channels were recorded at a 12000 Hz sampling frequency during the experiment. A multi-channel data acquisition recording system (system3, TDT, Alachua, FL) was used to capture microphone, feedback, electrical stimulation triggering TTL pulses, and ECoG signals with a common time scale.

### Vocal Response Times

Auditory voice onset and offset times were determined offline using custom Praat (Boersma and Weenik, 2018) and Matlab code. We report median (percentiles 25 and 75) as measure of central tendency, given that response times distributions were not normal (*See* *Figure2*), and our number of data points was small. Trials in which the subject started vocalizing more than 2 seconds after cued were considered no-responses. We excluded from vocal response analysis trials in which the subject started vocalizing before the cue display (n=1) and no-response trials.

### Voice F0

Voice pitch traces were estimated in absolute Hz, using a standard autocorrelation-based pitch tracking method implemented in Praat (Boersma, 1993) with a fixed 0.001 s step. Pitch traces were exported to Matlab for further analysis using custom code. Noisy pitch track segments and those with octave errors were identified and excluded. If the noisy segment was longer than 50% of the period of interest, the whole trial trace was excluded from the relevant analysis. For the evaluation of the pre-shift period we excluded from analysis and figures the initial 50 ms of F0 after vocalization onset.

### Peak F0 responses to pitch-shifted auditory feedback

Given that F0 dynamics could differ importantly between stimulation and non-stimulation trials before the pitch-shift was introduced, the selection of a baseline period affected dramatically the measured magnitude of a vocal response. We averaged the non-normalized traces (in Hz) separately in non-stimulation and stimulation trials per block. These average traces were detrended before identifying the most prominent peak in the response period (0 to 400 ms after pitch-shift onset) using the Matlab function *findpeaks*. We repeated this procedure with the inverted traces to identify potential negative peaks. We recorded the magnitude and polarity of the most prominent peak in the response period.

### Data availability

Data are available from the authors upon reasonable request.

## SUPPLEMENTARY FIGURES

**Supplementary Fig1.**
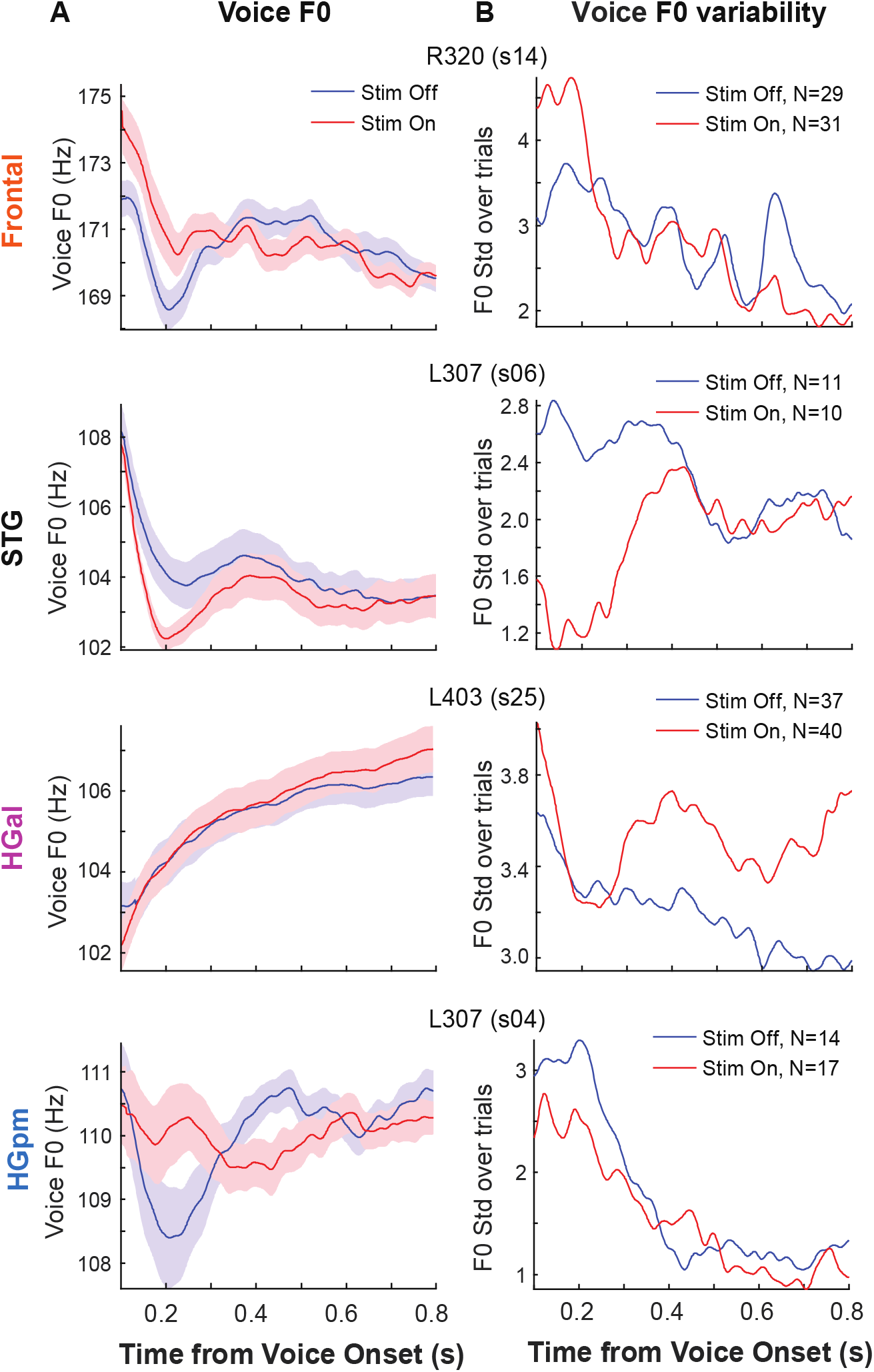
Voice F0 in the pre-shift period: Individual Block Examples. Column **A** shows average F0 traces (Hz) in stimulation (red) and non-stimulation (blue) utterances for one representative block per ROI. Shaded areas depict standard error over utterances. Column **B** shows the standard deviation over utterances in stimulation (red) and non-stimulation (blue) trials for the corresponding blocks.

**Supplementary Fig2.**
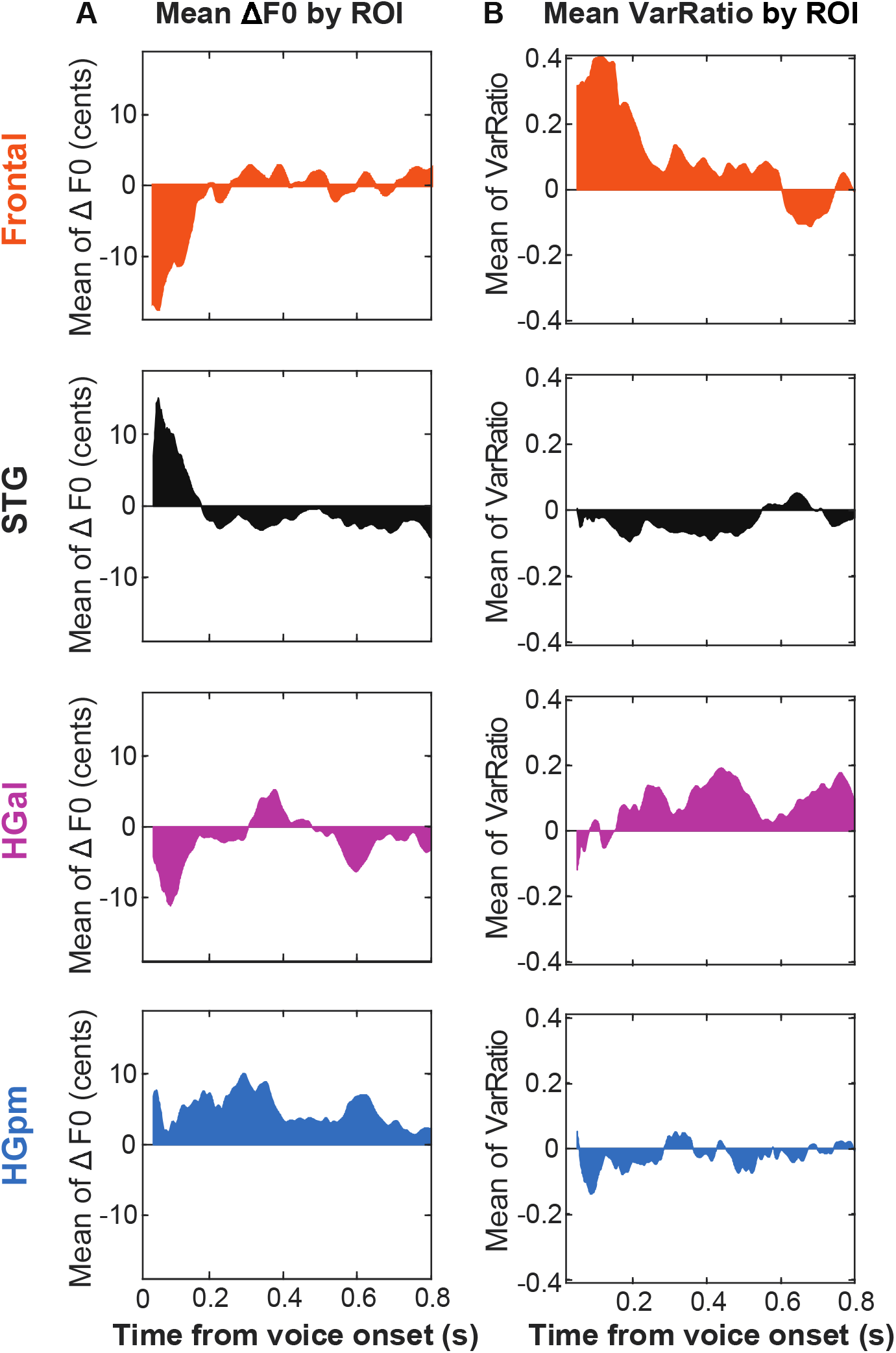
Voice F0 in the pre-shift period: Average functions per ROI. Column **A** shows the average DES-induced change per ROI after averaging across sites. Column **B** depicts average per ROI.

## Notes

### Competing Interest Statement

The authors have declared no competing interest.

